# What is particular about microplastics? A meta-analysis of the toxicity of microplastics and suspended sediments

**DOI:** 10.1101/2022.11.01.514507

**Authors:** M. Ogonowski, M. Wagner, B. Rogell, M. Haave, A. Lusher

## Abstract

Microplastics (MP) are perceived as a threat to aquatic ecosystems but bear many similarities to suspended sediments which are often considered less harmful. It is, therefore pertinent to determine if and to what extent MPs are different from other particles occurring in aquatic ecosystems in terms of their adverse effects. We applied meta-regressions to hazard data extracted from the literature and harmonized the data to construct Species Sensitivity Distributions (SSDs) for both types of particles. The results demonstrate that the average toxicity of MPs is approximately one order of magnitude higher than that of suspended solids. However, the estimates were associated with large uncertainties and did not provide very strong evidence. In part, this is due to the general lack of comparable experimental studies and dose-dependent point estimates. We, therefore, argue that a precautionary approach should be used and MP in the 1–1000 µm size range should be considered moderately more hazardous to aquatic organisms capable of ingesting such particles. Organisms inhabiting oligotrophic habitats like coral reefs and alpine lakes, with naturally low levels of non-food particles are likely more vulnerable, and it is reasonable to assume that MP pose a relatively higher risk to aquatic life in such habitats.

**Synopsis:** A meta-analysis indicates that microplastics are one order of magnitude more toxic than suspended sediments/solids, an estimate surrounded by considerable uncertainty.

**Graphical abstract:** 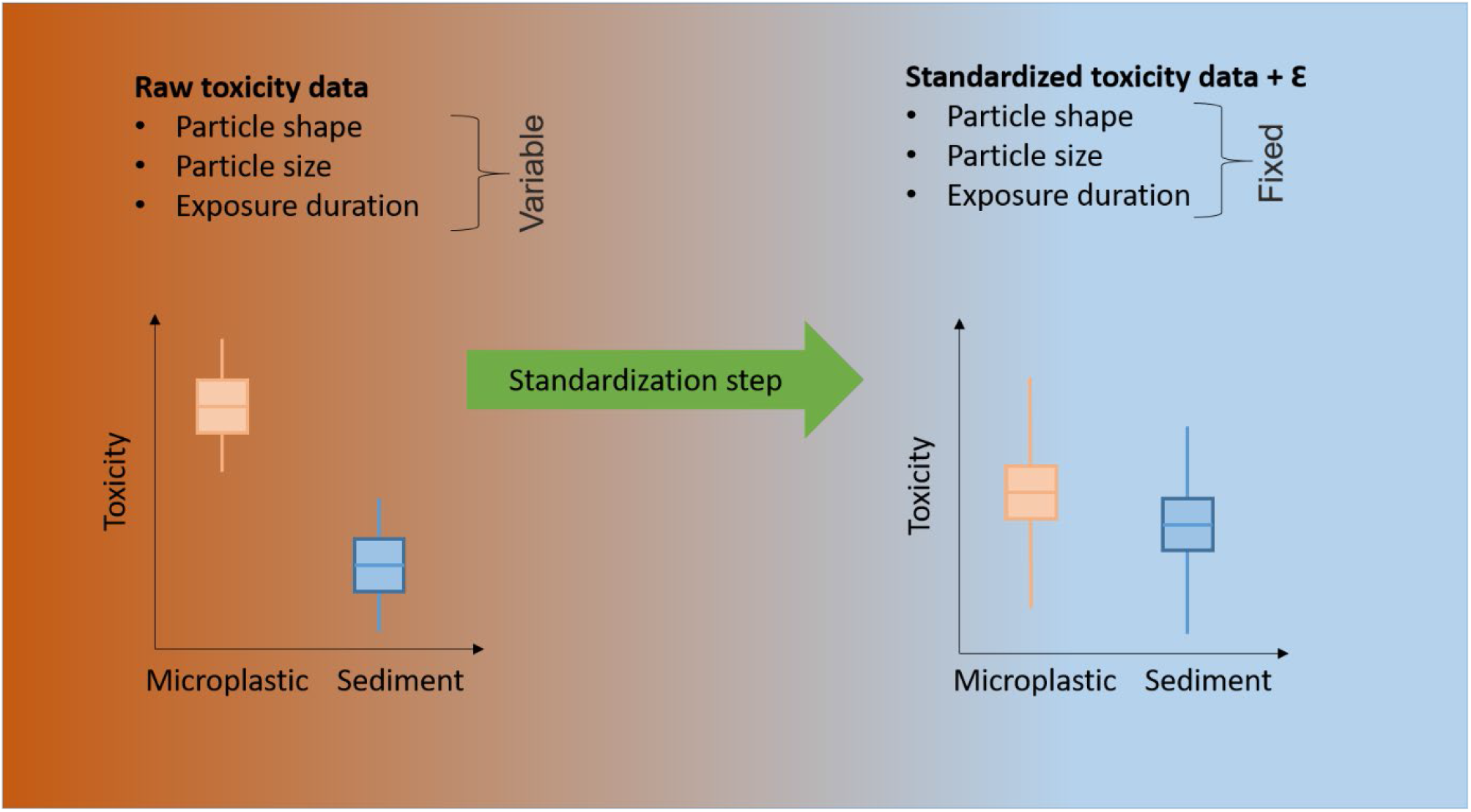

## Introduction

Microplastic pollution has emerged as a potential threat to the environment. This has spurred the development of a rapidly expanding research field aiming to quantify the hazard and risk of these pollutants. Assessments of both hazards and risks are complicated by (1) the heterogeneous nature of microplastics (MP), (2) the lack of standardized test methods (Redondo-Hasselerharm et al. 2018, Gerdes et al. 2019, Gouin et al. 2019), (3) the general difficulty in identifying and quantifying MP in complex environmental samples (Cowger et al. 2020, Lusher et al. 2020), and (4) the lack of data comparability driven by inconsistent reporting of MP characteristics (Cowger et al., 2020; Provencher et al., 2020). As a consequence, quantitative risk assessments of MP (Burns and Boxall 2018, Everaert et al. 2018, 2020, Adam et al. 2019, 2021, Besseling et al. 2019, Yang and Nowack 2020) have been criticized for the lack of alignment between exposure and hazard data (Koelmans et al. 2020). More specifically, the problem stems from mismatches between the size, shape and density of particles used in ecotoxicological test assays and those actually quantified in the environment.

Drawing inferences from such data is difficult, and Koelmans *et al*. (2020) proposed to overcome these issues by rescaling hazard and exposure data to a comparable distributions of particles. This method rests on the assumption that MP are inert particles and that the main mode of toxic action is food dilution, implying that the physicochemical properties of the particles are less important. Assuming that food dilution predominates as a major mechanism also implies that MP and any other non-food particles present in the environment are analogous with regards to their effects. In fact, naturally occurring, non-palatable particles like suspended sediments (SS), chitin and cellulose are known to induce similar effects in aquatic organisms (Newcombe and Macdonald 1991, Gordon and Palmer 2015, Ogonowski et al. 2018). Consequently, this begs the question whether MP are unique with regards to their toxicity, or whether they are toxicologically identical to other non-food particles. This question is important because, depending on the answer, MP would either need to be studied, assessed, and managed as a specific group of contaminants or be considered as an integral component of suspended matter. It is, therefore, important to determine if and to what extent MP differ from other particulate matter present in the environment.

A wide range of experimental conditions has been used for testing the toxicity of anthropogenic particles to aquatic organisms. This has resulted in a high level of heterogeneity in exposure conditions and experimental designs that, on the one hand, provides insight into the likely effects of various exposure scenarios. On the other hand, it hampers comparability across studies. The variability in test conditions means that any analysis of literature data aiming to assess the relative toxicity between different particulate stressors needs to be able to account for several factors pertinent to (1) the test materials, (2) the sensitivity of the test species and specific endpoints, and (3) the experimental conditions (e.g., exposure duration).

The most straightforward approach to achieve comparable data is to subset data to one or several common denominators. However, this approach removes valuable information and is rarely feasible in practice due to the scarcity of studies with comparable experimental designs. One way to solve this misalignment is to statistically account for the variability and normalize the data to a common scale, which can be achieved using various multiple regression techniques (Thompson and Higgins 2002, Sun et al. 2021). For example, if the goal is to compare the toxicities between different materials that also differ in particle size (a characteristic known to affect toxicity), we can statistically control for the difference in particle size. This approach will give a comparable test of the toxicity of the materials that is independent of particle size.

Here, we address the question of whether MP are toxicologically different from naturally occurring particles by evaluating the relative toxicity of plastic particles and suspended sediments in the size range 1-1000 µm based on a comprehensive set of published ecotoxicological data. Using a series of probabilistic approaches coupled with data standardization, we account for the uncertainty in the data while keeping factors influential to the biological responses at fixed levels. This allows us to obtain more comparable measures of toxicity and, consequently, perform an improved hazard assessment.

## Materials and Methods

### Literature review and compilation of ecotoxicity data for microplastics and suspended sediments

Hazard data from toxicity studies with MP were collected by means of a systematic review, covering the period January 2016 - February 2019 published by the Norwegian Scientific Committee for Food and Environment (VKM 2019). Details regarding the search criteria and the selection process are provided in Appendix I of the VKM report, and the raw data is provided as Supporting Information data table 1.

Data collection for toxicity studies with suspended sediments were mainly based on studies from previous data compilations and reviews (Gordon and Palmer 2015, Ogonowski et al. 2018) but complemented with additional searches using Web of Science, Scopus and Google Scholar using the following search terms: “suspended solids”, “suspended matter”, “suspended material”, “sediment”, “mineral particles”, “effects”, “aquatic”, “filter feeder” alone or in combination. In contrast to the search for MP hazard data, no restrictions on publication date were made. For some older studies listed by Gordon & Palmer (2015), the original manuscripts were unavailable. In those cases we used the toxicity data as reported in the paper by Gordon & Palmer (2015).

Since manual data extraction for systematic reviews is prone to errors (Mathes et al. 2017), we conducted an additional error screening after the dataset had been compiled. Twenty percent of the data entries (rows in the dataset, Supporting Information data table 1) were selected at random to be reassessed by three of the co-authors. We found minor errors relevant to the data analyses in 6 out of 40 endpoints. Out of these six errors, five occurred in one publication. As we account for publication ID in the analysis we conclude that the risk of systematic errors is small and unlikely to affect our conclusions.

The particle size, either reported as the mean/median size or by visual inspection of the actual size distribution in each study was assigned to sediment grain size classes according to the Wentworth scale (Wentworth 1922). Size distributions that spanned several grain size classes were assigned to the most predominant class. The division into size classes was necessary as some studies, in particular the SS studies, lacked clearly defined size distributions. For reasons of consistency, we used a nominal particle density for the material used in the studies (Supporting Information data table 1).

The primary aim of the literature search was to compile a dataset for hazard assessment. For this purpose, we extracted effect concentrations reported in each study in the form of the lowest observed effect concentrations (LOEC), effect concentrations derived from dose-response relationships (EC_10_, EC_20_, EC_50,_ LD_50_), and no-effect concentrations defined as the highest observed-no-effect concentration (HONEC). The raw toxicity data in the form of varying dose descriptors other than the no-observed effect concentrations (NOECs) were converted to estimated NOECs using a conversion factor specific to each descriptor (Adam et al. 2019).

### Data subsetting to make datasets comparable for hazard assessment

The compiled dataset was restricted to studies in which aquatic organisms were directly exposed to MP or SS added to the medium. Thus, studies in which the particles were incorporated into food or delivery via trophic transfer were excluded. Data on fibrous particles were omitted since this particle shape was exclusive to MP. Studies employing particle sizes < 0.98 µm (clay-sized particles) were also removed since the mode of toxic action for nano-sized particles may be different due to their capacity to pass biological barriers and cell membranes (Matthews et al. 2021). Since the main mode of action of microparticles > 1 µm is assumed to be food dilution (de Ruijter et al. 2020), we further restricted the data to only contain test organisms where the main route of exposure was through ingestion. This also excluded toxicity data involving primary producers, non-feeding larval stages, eggs, and embryos. We only considered higher levels of biological organization (Galloway et al. 2017), i.e. individual and population level endpoints limited to *growth, mortality* and *reproduction* since lower-level endpoints may represent transient responses. The subsetted data used for analysis consisted of 43 studies (MP = 28, SS = 16) and 200 biological endpoints (MP = 123, SS = 77) and is provided as supplemental material (Supporting Information data table 1).

### Conversion from numerical and mass-based concentrations to volumetric units

The choice of dose metric, such as volume, mass, or number of particles depends on a toxicant’s main mode of action. The appropriateness of a particular dose metric for solid substances is still under debate, particularly in the field of nanomaterial toxicology (Delmaar et al. 2015, Teunenbroek et al. 2017). Assuming that MP as well as SS mainly affect organisms by means of food dilution, the correct dose metric should be based on volume of particles per volume of medium.

For studies where spherical particles were used and effect concentrations were reported as particle numbers per volume, we firstly converted the numerical concentrations to a mass-based concentration according to the following equation:

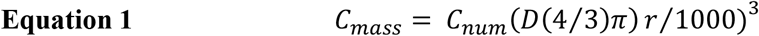

Secondly, the mass-based concentrations were converted to volumetric concentrations as

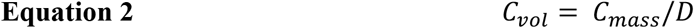

Where C_mass_ = the mass-based concentration (mg L^-1^), C_num_ = the numerical concentration (number of particles L^-1^), D = polymer density (g cm^-3^), r = the particle radius in µm and C_vol_ = the volumetric concentration (mm^3^ L^-1^). This conversion was necessary to explicitly account for differences in particle density between studies.

### Probabilistic modeling of the relative toxicity of microplastics and suspended sediments

We used two slightly different statistical models to compare the relative toxicity of MP and SS to assess the robustness of the predicted hazard. Both methods have a probabilistic foundation as to account for the uncertainties in the data but differ somewhat conceptually. Since MP are considered emerging pollutants, we wanted our results to be conservative in terms of not providing a false negative conclusion. We hence used an alpha level of 10% rather than the conventional 5% when comparing toxicities between MP and SS.

### The hierarchical standardized pSSD+ model

To compare the hazards across species, we followed the probabilistic SSD-approach first proposed by Gottschalk & Nowak (2013) and more recently adopted by Adam et al. (2019, 2021) for the risk assessment of MP. In brief, the pSSD+ model developed by Adam and colleagues does not assume the data to fit any specific theoretical distribution and avoids the loss of valuable data by incorporating all available toxicity data at the species level instead of using mean estimates.

In the pSSD+ model, the uncertainties in the underlying data are accounted for using arbitrary uncertainty factors. Instead of using such an *ad-hoc* approach, the heterogeneity can instead be modeled from the data using well-established multiple regression techniques (Thompson and Higgins 2002, Sun et al. 2021). Such an approach also has the advantage of estimating the toxicity for specific particle sizes, shapes, exposure times and other parameters. Thus, we utilized a two-step hierarhical approach to model the SSDs. In the first step, we used a Bayesian regression as implemented in the R-package *brms* (Bürkner 2017) to predict the toxicity of MP and SS for a fixed set of parameters based on the collated literature data. In other words, we estimated the toxicity for each particle type separately, while keeping particle size, shape and exposure duration constant, making data comparable. The probabilistic model also enabled the uncertainty in the estimated toxicity to propagate through all analytical steps. The precursory standardization model can be described as a basic linear regression model:

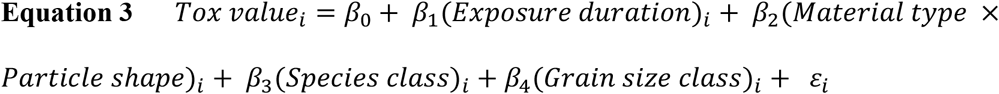

The toxicity data, translated into volumetric concentrations [mm^3^ L^-1^] (Koelmans et al. 2020) was standardized, by fixing the *Exposure duration* to 28 days, which was the upper threshold in our data for a chronic exposure (Table S2). *Particle shape* was set to “irregular” since this is a shape that encompasses both MP and natural suspended solid particles. *Material type* and *Particle shape* were modeled as a composite categorical factor (equivalent to an incomplete factorial interaction term) with three levels (SS-irregular, MP-irregular and MP-spherical) since no spherical SS particles were included in our dataset. *Grain size class* was set to “clay” (0.98–3.9 µm), a common size category for the entire dataset. In contrast to standard SSDs (Kooijman 1987, Aldenberg and Slob 1993), we grouped species-level data to taxonomic class level instead, in order to not overparameterize the model. This was motivated by the assumption that closely related species have similar feeding modes and sensitivities to particle exposures. Specification of the priors is provided in the Supporting Information and Table S3.

In the second step, the predicted and standardized toxicity values were used as input to the pSSD+ model described by Adam et al. (2019, 2021) to produce two SSDs, one for MP and another for SS. To retain the uncertainty in the toxicity estimate throughout the analytical process, we used the 5^th^ and 95^th^ percentile of the posterior distribution for each predicted toxicity value as input in the pSSD+ model. The relative hazard of MPs and suspended sediments was evaluated by comparison of the 5^th^ percentile-hazardous concentration (HC_5_) from the two standardized pSSD+ models and by comparing the full posterior distributions of the HC_5_-values. Following the studies by Adam *et al*. (2019, 2021), HC_5_ was considered equal to the predicted no-effect concentration (PNEC). The pSSD+ model was generated using 10,000 random permutations.

### Alternative approach to compare the hazard of microplastic and suspended sediments

In order to validate our approach, we also analyzed our data in an alternative framework using a Bayesian mixed model. The model was used to predict a NOEC (pNOEC) for MP and SS while accounting for the variability in experimental conditions, particle characteristics, exposure duration and variation across taxonomic groups and studies. In contrast to the pSSD+ model, this model accounted for the fact that no-effect studies (HONEC) are right-censored and studies reporting LOECs are left-censored when the LOEC equals the lowest test concentration by explicitly incorporating this uncertainty into the model. Hence, the intention was twofold: (1) to compare chronic NOEC posterior distributions (the relative toxicity) between MPs and suspended sediments while statistically controlling for other explanatory variables, and (2) to identify potential drivers of the toxicity.

The criteria and statistical approaches for determining hazardous or safe concentrations differ among studies which makes toxicity data not directly comparable. To align toxicity data to the same scale, it is common to apply uncertainty factors (UF) to derive the chronic NOEC. Although there is no consensus on what these UFs should be, Wigger *et al*. (2020) suggested a range of conversion factors; one for the dose descriptor conversion (UF_dose_, Table S4) and another one to convert acute to chronic toxicity data (UF_time_). Along these lines, we used the estimated NOEC (*eNOEC*) as the response variable in the model calculated by dividing the reported dose descriptors by the appropriate uncertainty factor (UF_dose_). Contrary to the common approach (Adam et al. 2019, 2021, Wigger et al. 2020), we did not multiply UF_dose_ with UF_time_ to derive the chronic NOEC. Instead, we modeled the effect of exposure duration as a predictor in the model. By doing so, we estimated the effect directly from the data instead of using an arbitrary UF with an *ad hoc* associated uncertainty, thus avoiding the assumption of a positive relationship between exposure time and toxicity a priori.

Toxicity values that equal the highest or the lowest employed test concentrations are so called “censored” data. This means that the true hazardous concentration exceeds the tested concentration range and is unknown. The censoring of HONEC and LOEC-values was accommodated using the *cens-*function in the R-package *brms*. Apart from the focal variable *Material type*, the variables *Feeding strategy, Particle exposure, Grain size class* and *Particle shape* were included as co-variates in the model because they are intimately linked to exposure, food processing and the organism’s sensitivity to particles. To account for the fact that *Particle shape* and *Material Type* were not fully crossed (spherical shape missing from the suspended sediments dataset), we modeled the interaction of these two variables as a single composite factor (the fused combination of *Particle shape* and *Material Type*) the same way as for the precursory pSSD+ model. *Grain size class* was moreover modeled as a monotonic variable due to the ordered nature of the size classes (Bürkner and Charpentier 2015). To account for the variability between studies, we considered *Study* as a random effect on the intercept. To account for variability across species we initially also included *Species* as a random intercept term together with the interaction between *Species* and *Study*, but this resulted in a too complex model and unsatisfactory low effective sample size for the interaction term. The final model thus omitted the *Species* term but retained the interaction term which rests on the assumption that the sensitivities of species within a particular study are more similar than across studies (Table S2). The error structure was modeled as a t-distribution to accommodate the presence of outlying data points further out in the tails of the distribution. For all models, we ran four chains with 5000 iterations each after a burn-in of 2500 iterations was discarded. Thinning was set to one. Markov chain Monte Carlo (MCMC) convergence to the equilibrium distribution was monitored visually using the *bayesplot* package (Gabry et al. 2019) and by evaluation of 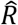 values and effective sample sizes. We found no sign of failed convergence and 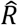-vales were equal to one, indicating that the MCMC chains had converged at similar values. Model residuals for the Bayesian mixed model were evaluated visually (Figure S1). Specification of the priors is provided in the Supporting Information and Table S5.

## Results and Discussion

### Microplastic toxicity compared to suspended sediments

Based on the Bayesian mixed model, the standardized mean NOEC for MP on the data scale was approximately 12-fold lower compared to that of SS but the credible interval for the groups overlapped, indicating no significant difference between MP and SS (Table S1). However, the one-sided 90% probability distribution of the difference in posteriors did not contain zero, suggesting that MP are more hazardous than SS. The mode of the difference in posterior distributions was centered around one unit on the log10-scale, corresponding to a ten-fold difference in toxicity (Figure 1). This pattern was consistent with the standardized pSSD+ model where the PNEC-distributions for MP and SS overlapped (Figure 2) but the one-sided 90% probability distribution of the difference in PNEC posteriors did not overlap zero (Figure 3) and the PNEC for MP was 7.7 times lower than that of SS, suggesting that species likely are more sensitive to MP exposure compared to other particulate matter. This is also in line with a previous assessment based on a smaller dataset where the LOEC (at individual and population level) was significantly lower for MPs compared to suspended sediments (Ogonowski et al. 2018). It is, however, important to consider that the differences in hazard can be attributed to other causes than actual differences in toxicity, such as differences in experimental designs and exposure conditions that are difficult to account for statistically.

**Figure 1.**
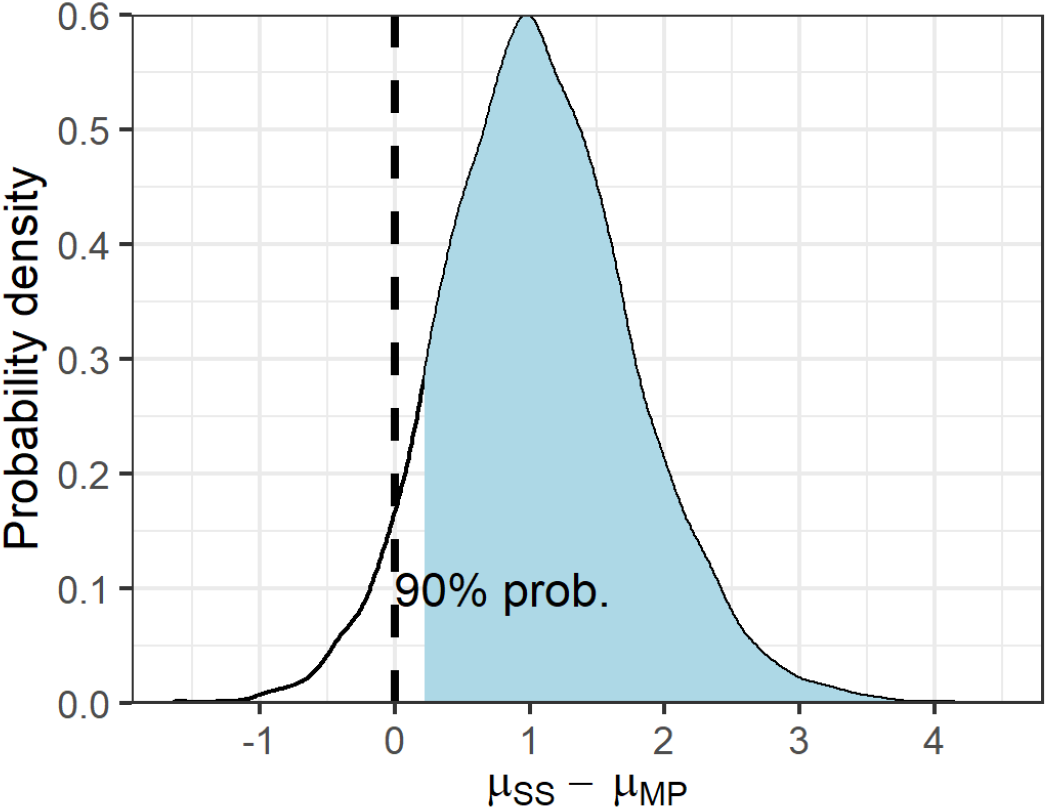
Marginal mean difference in posterior probabilities between suspended sediments (SS) and microplastic (MP) groups in the censored, Bayesian mixed model (model 2, Table S2). The shaded area shows the 90% probability (one sided test) for MP to have a lower eNOEC compared to SS.

**Figure 2.**
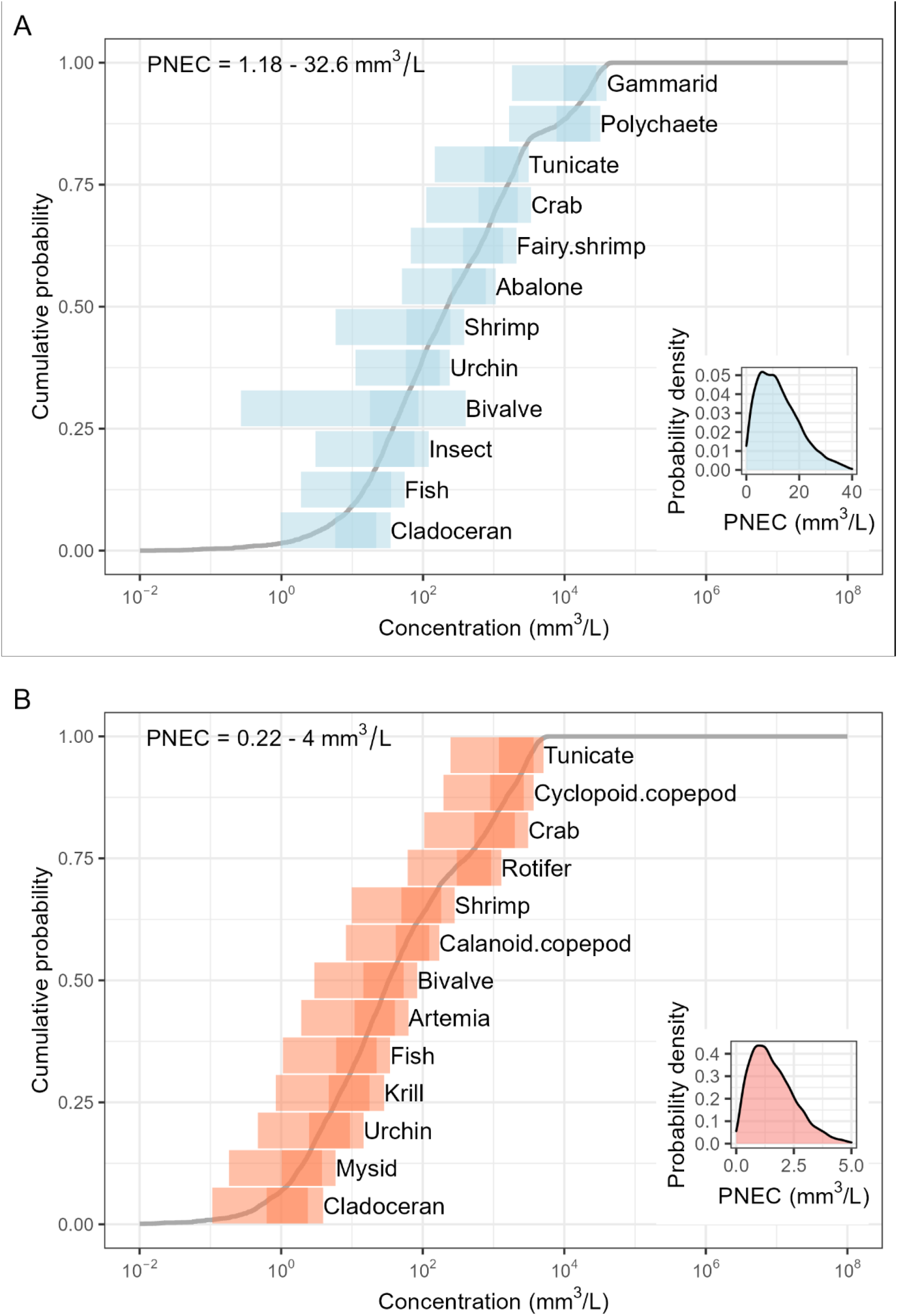
Probabilistic species sensitivity distribution based on volume-based toxicity data corrected for inter-study differences in particle characteristics and exposure conditions; A) for suspended sediments and B) for microplastics. The dark shaded horizontal bars represent the 25-75^th^ percentile ranges and lighter shaded area the 5-95^th^ percentile range.

**Figure 3.**
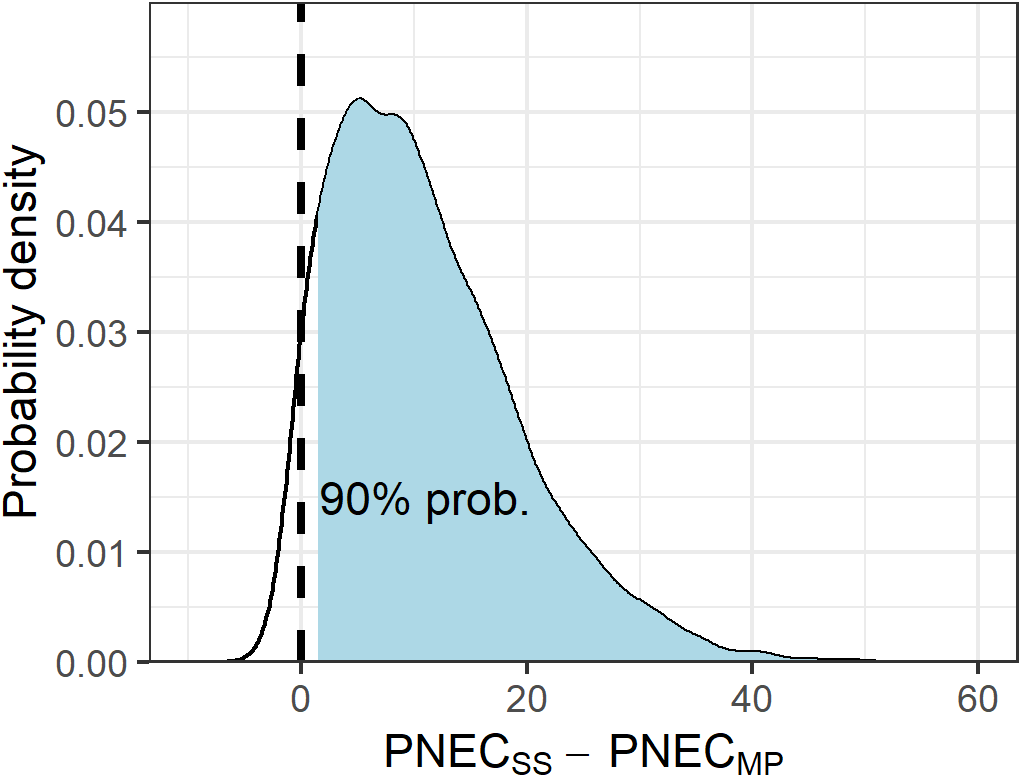
Posterior distribution of the difference in PNEC values (mm^3^ L^-1^) between suspended sediments (SS) and microplastic (MP) SSDs. The shaded area shows the 90% probability (one-sided test) for MP to have a lower PNEC compared to SS.

### The use of model particles in test assays yields unrealistic toxicity estimates

Two aspects, we did not capture statistically, may result in a higher toxicity of MP compared to SS. Both relate to the use of pristine MP versus SS in toxicity studies. First, MP will leach plastic chemicals that can, at least in some cases, drive the overall MP toxicity (Martínez-Gómez et al. 2017, Heinrich et al. 2020, Zimmermann et al. 2020, Beiras et al. 2021). In addition, commercially available MP can contain preservatives (e.g., sodium azide) that exacerbate the particles’ toxicity (Yang and Nowack 2020). Accordingly, the hazard data we used here may include the toxicity of plastic chemicals and preservatives, which does not occur in studies with natural particles. Like MP, SS can also contain chemicals adsorbed from their environment, such as polycyclic aromatic hydrocarbons, polychlorinated biphenyls (PCBs) and other persistent pollutants (Santiago et al. 1993, Rügner et al. 2019) Thus, SS toxicity may also be caused by their physical and chemical composition (Rivetti et al. 2015, Lu et al. 2021). However, since many of our SS-studies used pristine, unconditioned mineral particles (78% of the SS endpoints, SI data table 1) they likely underestimate the toxicity that would occur under natural conditions. Conversely, the presence of toxic preservatives and other plastic chemicals such as UV stabilizers, surfactants and monomer residues which are specific to some MP studies can leach from the MP during exposure and induce chemical toxicity. Indeed, Yang and Nowak (2020) demonstrated for nanoplastics that removing hazard data for particles that contained sodium azide resulted in higher PNECs (i.e., lower toxicity).

To compare MP and SS particles on equal terms, the potential effects of leachable chemicals should be accounted for either by removing data for particles containing chemicals (e.g., if they have not been washed) from the meta-analysis or by modeling it statistically (e.g., as covariates in the meta-regression). However, to do so completely and without bias would be practically impossible because the presence of reported toxic preservatives likely is non-random and skewed towards commercially available MP. In addition, the latter may also contain a multitude of other proprietary and undisclosed chemicals which cannot be easily accounted for (Heinrich et al. 2020). Hence, we chose to treat the chemical component as an integral part of the toxic response, not discriminating between specific physical and chemical toxicity. This results in a higher-than-expected average toxicity but also higher uncertainty of that estimate.

A second aspect regards the fact that MP which are aged under natural conditions, will be more comparable to SS and have a different toxicity than pristine MPs which dominate our dataset. This has been demonstrated experimentally with some studies reporting a lower toxicity of aged vs pristine MP (Zou et al. 2020, Schür et al. 2021) and other studies reporting an increase in toxicity after weathering (Zhang et al. 2021, 2022). Although the causes of the altered toxicity are not fully understood, they seem to be related to the disassociation of plastic chemicals, the formation of a protein corona and biofilms as well as the fragmentation into smaller nano-sized particles. The latter can increase toxicity of the overall particle mixture during ageing but may reduce the toxicity of the particles in the same size fraction as pristine MP (Zhang et al. 2021, 2022).

The sorption of biomolecules on the particle surface (eco-corona) and ultimately biofilm formation (Galloway et al. 2017) may promote particle aggregation and larger average particle size (Porter et al. 2018, Michels et al. 2018, Motiei et al. 2021). Although the same would be theoretically true for mineral particles there is evidence to suggest that these particles do not aggregate to the same extent as MP (Motiei et al. 2021). This shift in particle sizes may lead to the formation of MP aggregates that are too large to be consumed, which may decrease their bioavailability and hence their capacity to cause adverse effects. In addition, the eco-corona or biofilm on particles can provide extra nutrition and, thus, counteract food dilution effects for some types of aging (incubation in nutrient rich raw wastewater) but not for others (e.g., treated wastewater and river water which are lower in nutrients and microbial activity) (Amariei et al. 2022). Whether and to which extent such modulation also applies to SS is not clear from the literature. Given the dearth of studies on the toxicity of aged MP, we could not account for this factor in our meta-analysis. Consequently, our MP and SS toxicity estimates are likely not directly translatable to natural systems since they reflect somewhat artificial conditions.

### Differences in test-concentration ranges affect the predicted hazard

The pSSD+ model does not account for the fact that no-effect studies are right-censored (undefined upper effect concentration) or that LOEC values can be left-censored if they equal the lowest used test concentration (undefined lower effect concentration). This may lead to an over- or underestimation in toxicity, respectively. In our data collection, the distribution of no-effect data was unequal across MP and SS studies with a higher frequency of such data points in the MP data compared to the SS data (69.9 vs. 24.7%, Table S6). Also, neither of our models accounts for the fact that the experiments were conducted using different concentration ranges. Experiments involving natural suspended solids or minerals usually employ test concentrations in the order of grams L^-1^ (Cohen et al. 2014) to cover a natural range of concentrations. MP studies on the other hand use orders of magnitude lower concentrations, due to the desire to test “environmentally relevant” concentrations. In fact, in our data, the average highest concentration for SS was two orders of magnitude higher compared to the MP studies (Table S6). Although many MP studies have been criticized for using unrealistically high test concentrations (Lenz et al. 2016, Connors et al. 2017, Cunningham and Sigwart 2019), these concentrations are still much lower than naturally occurring levels of SS. The difference in concentration ranges poses a problem when the objective is to compare the hazard of different toxicants based on dose metrics like the LOEC or the NOEC – which are directly dependent on the range of test concentrations used (Laskowski 1995, Warne and Van Dam 2008, Fox and Landis 2016, see also tables S4 and S6). The high proportion of no-effect studies in the MP data indicates that the hazardous concentrations likely are higher and closer to those of SS than our statistical models suggest.

Even though an uncertainty factor was applied to adjust for the unknown effect concentration it may not have been large enough. Such disparities in the experimental designs cannot be statistically accounted for unless dose dependent point estimates are used exclusively (Van Der Hoeven et al. 1997). This was, however, not possible due to the general lack of such data, in particular for MP. Excluding HONEC-data from the model was on the other hand not feasible either because it resulted in a too complex model for the data. Moreover, the removal of censored data (e.g., HONEC) usually results in biased estimates and variances (Turkson et al. 2021, Bouaziz n.d.).

The best approach to make data fully compatible would be to perform paired comparisons of different particle types within an experiment where the exposure conditions are the same. The use of natural reference particles in MP testing has recently been advocated (Ogonowski et al. 2016, 2018, Connors et al. 2017, Scherer et al. 2018, Gerdes et al. 2019, Gouin et al. 2019, Arp et al. 2021) but it’s adoption has until recently been comparatively scarce in the scientific literature. We argue that the use of reference particles such as natural minerals is a way to increase the ecological relevance of ecotoxicological studies since it provides a benchmark for particle toxicity (Scherer et al. 2020, Schür et al. 2020). Such setups will also help to identify the mechanisms driving toxic responses to particles.

### Drivers of toxicity

The Bayesian mixed model enabled the toxicity data to be standardized and comparable between MP and SS studies. The heterogeneity in the data was large and varied considerably between specific studies (e.g., experimental setups) and test species. The proportion of variance explained by the variability across studies was on average 81% and the 95% Credible Interval (CI) ranged 33 – 84%. Variability across species nested within studies was on the other hand lower (24%, 95% CI = 11 – 46%). Although this high level of variation is expected given the wide variety of test materials, species and experimental designs, this contributed to a high degree of uncertainty in the regression coefficients, with the credible intervals all overlapping or being close to overlap, indicating a low degree of confidence (Table S1, Figure S2). In this context, one advantage of Bayesian over frequentist models is their ability to make probabilistic statements regarding the parameter estimates, which allows for a more nuanced interpretation. A closer inspection of the central tendencies of the coefficient posteriors reveals that even though the overall uncertainty was high, the highest probability densities were centered away from zero for several variables (Figure S2, Figure 4). Notably, the probability of *Grain size class* to have a positive slope (one-sided evidence ratio) was > 95% suggesting decreasing toxicity (higher pNOEC) with increasing particle size. This is in line with previous observations of MPs in the current size range (Ziajahromi et al. 2018). We can also see that 39% of the total change in pNOEC due to *Grain size class* happens between the first two predictor categories (i.e., clay and silt, Table S1) which indicates that the relationship is non-linear. It is probable that very fine particles have additional effect mechanisms apart from food dilution, such as an obstruction of gas exchange through the gills in fishes and invertebrates (Hess et al. 2015, 2017, Lowe et al. 2015, Watts et al. 2016), clogged feeding appendages in filtrating invertebrates (Cole et al. 2013, Savinelli et al. 2020) or tissue translocation with potential consecutive down-stream effects (Haave et al. 2021).

**Figure 4.**
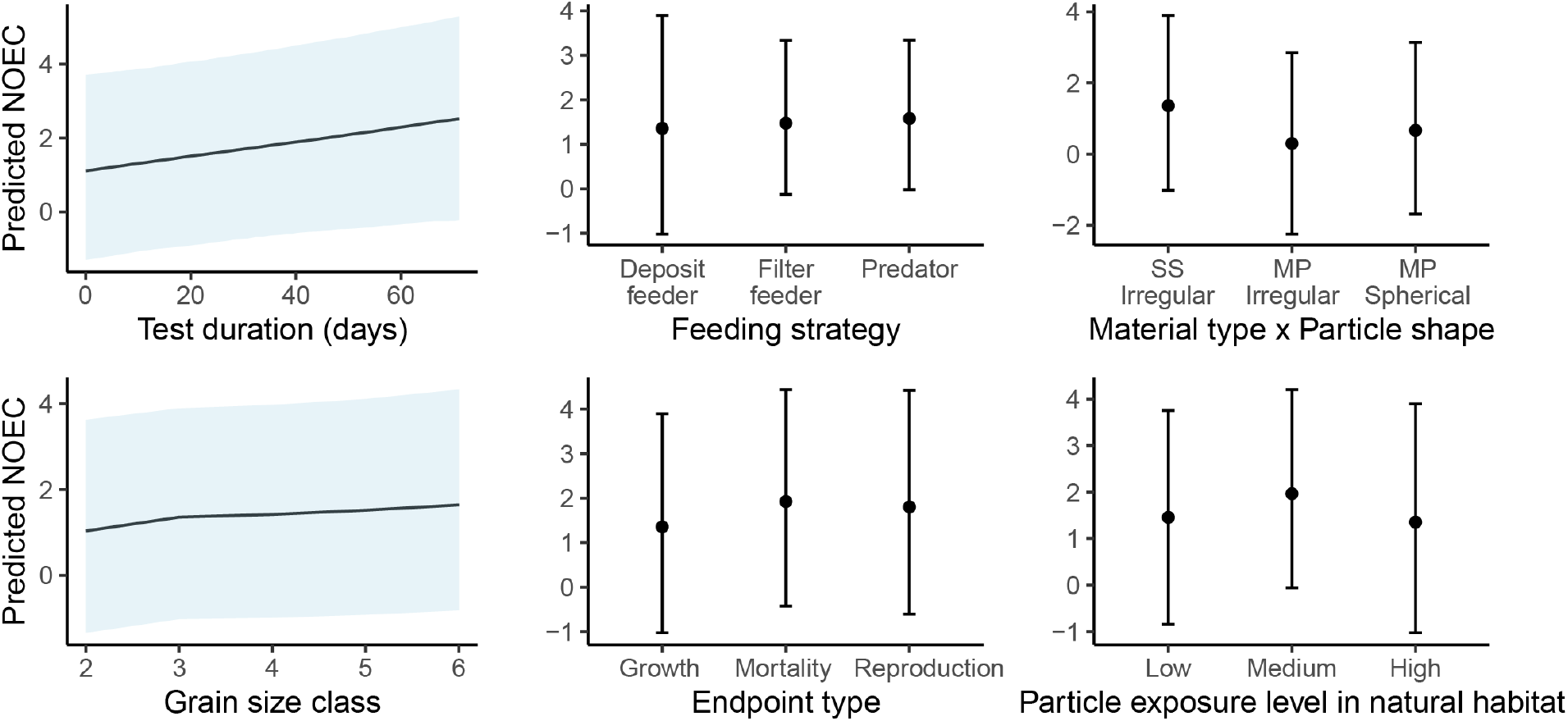
Marginal means plot for the explanatory variables in the censored, Bayesian mixed model (model 2, Table S2).

As for the variable *Grain size class*, we saw the same pattern for *Exposure duration* (Figure 4). Although decreasing toxicity with increasing exposure time may seem counterintuitive at first, it is plausible in circumstances where sedimentation is allowed to occur without renewal of the test medium or an effect of increased food intake due to the secondary ingestion of nutritious biofilms associated with the particles (Amariei et al. 2022). Alternatively, it can be an artefact linked to the fact that experiments with longer exposures tend to employ lower test concentrations which is problematic when concentration dependent dose metrics, like LOECs and NOECs, are used (Supporting information Figure S3-Figure S6). The failure to control for such effects can bias toxicity assessments when particles of different density and sedimentation rates are compared, in particular for suspension feeding organisms (Connors et al. 2017, Ogonowski et al. 2018, Gerdes et al. 2019, Gouin et al. 2019). Although such experimental designs have been rather common in the past, procedures to overcome these shortcomings have recently been proposed (Gerdes et al. 2019, Motiei et al. 2021). Albeit not fully conclusive, the pattern of decreasing toxicity with exposure time remains when the more robust dose-dependent point estimates (EC_50_) are considered (Figure S7) suggesting this is a true effect.

Moreover, we expected the sensitivity of a species/life stage to be linked to its native environment, meaning that organisms during different stages of development should be well adapted to cope with local turbidity levels (McFarland and Peddicord 1980). Contrary to our expectation, we did not find any coherent evidence to support this hypothesis. Species in the low turbidity category (variable particle exposure) had only a 45% probability to be more sensitive to particle exposures than the ones in the high turbidity category (one-sided posterior probability). However, compared to the intermediate turbidity, the probability was 86%, suggesting rather strong evidence for species adapted to a low-particle environment to be more sensitive than species that more often are exposed to higher levels of SS. We expected species classified to the high turbidity category to be the least sensitive, but we found low support for this hypothesis and the probability for the intermediate class to be more sensitive than the high turbidity class was merely 21%. One reason could be that our classification into the “low turbidity” group is more accurate as the boundary between clearwater and more tolerant species is simply more distinct and easily defined. Clearwater species were almost exclusively those inhabiting tropical and oligotrophic waters or species with a fully pelagic life history where the occurrence of suspended particles is naturally low (Supporting information data table 1). In contrast, it is more difficult to distinguish between the medium and high categories because most of these species and life stages are mobile and can switch between clear and turbid water and their exposure is generally dependent on seasonality and habitat specific conditions.

Out of the three selected endpoints (growth, reproduction and mortality), growth was the most sensitive and mortality the least (Figure 4, Table S1, Table S2, Figure S2). The higher sensitivity of the sublethal endpoints was expected and indicates that the model behaved as predicted. This also supports the hypothesis that food dilution and/or increased energy expenditure are important mechanisms when exposed to non-caloric particles because they compromise growth which in turn decreases reproductive capacity and ultimately leads to starvation (Madon et al. 1998, Wright et al. 2013, Ogonowski et al. 2016, Foley et al. 2018). It does however not exclude other possible modes of action for which Dynamic Energy Budget models or other individual-based modelling approaches would be needed.

Using a meta-regression based approach combined with a novel standardization step to harmonize data for SSD analysis, we have demonstrated that the average toxicity of MPs is approximately one order of magnitude higher than that of SS. However, the uncertainties around these estimates are large and the apparent difference in toxicity is partly due to systematic differences in experimental designs that cannot be accounted for statistically. Well-designed comparative experiments with plastic and non-plastic particles, where the potential effect of associated chemicals is accounted for and dose-dependent point estimates are derived, are needed to accurately assess the effects of MPs in the field relative to those of SS. In lack of better evidence, it is advisable to apply a precautionary approach. Hence, MP in the 1–1000 µm size range should, for the time being, be considered as moderately more hazardous to aquatic organisms capable of ingesting particles in this size range. Organisms inhabiting oligotrophic habitats like coral reefs and alpine lakes, with naturally low levels of non-food particles are likely more vulnerable, and it is reasonable to assume that MP pose a relatively higher risk to aquatic life in those areas.

## Acknowledgement

This work was funded by the Norwegian Scientific Committee for Food and Environment (VKM) and The Swedish Environmental Protection Agency (Naturvårdsverket) for the MIxT project [grant number 802-0160-18]. MW acknowledges funding from the European Union’s Horizon 2020 research and innovation programme under the Marie Sklodowska-Curie grant agreement No 860720 and the Norwegian Research Council (301157). We would also like to thank Prof. Elena Gorokhova, Department of Environmental Science, Stockholm University, Sweden for statistical advice and valuable comments on the manuscript.

## Potential competing interests

Martin Wagner is an unremunerated member of the Scientific Advisory Board of the Food Packaging Forum (FPF). He has received travel funding from FPF to attend its annual board meetings and from Hold Norge Rent (Keep Norway Beautiful) to speak at one of their conferences. Martin Ogonowski has received travel funding to present his research at the chemicals industry’s annual meeting in 2019 organized by IKEM – Innovation and Chemical Industries in Sweden and KTF – The Chemical Technical Industries (Kemisk Tekniska Företagen). All authors have received financial compensation from the Norwegian Scientific Committee for Food and Environment (VKM) for working on this dataset.

## Abbreviations

pNOEC: predicted No Effect Concentration
eNOEC: estimated No Effect Concentration
UF: Uncertainty factor
MP: microplastic
SS: suspended sediments
LOEC: Lowest Observed Effect Concentration
HONEC: Highest Observed No Effect Concentration
PNEC: Predicted No Effect Concentration

## Notes

### Summary of Updates

Supplementary data table 1 has been uploaded. This data table contains the raw data used in the analyses.

