## Supporting information for "What is particular about microplastics? A meta-analysis of the toxicity of microplastics and suspended sediments"

Table S1. Summary statistics from the Bayesian mixed model (table S2, model 2).

| *Response* | *log10(eNOEC)\| cens(censored)* | | |
| --- | --- | --- | --- |
| *Predictors* | *Estimates* | *std.error* | *CI (95% )* |
| (Intercept) | 0.81 | 1.34 | -1.72 – 3.43 |
| Particle exposure -High | *Reference* |  |  |
| Particle exposure -Low | 0.09 | 0.94 | -1.81 – 1.89 |
| Particle.exposure -Medium | 0.64 | 0.82 | -1.01 – 2.22 |
| Feeding strategy -Deposit feeder | *Reference* |  |  |
| Feeding strategy -Filter feeder | 0.13 | 1.04 | -1.89 – 2.13 |
| Feeding strategy -Predator | 0.22 | 1.03 | -1.72 – 2.19 |
| Endpoint category -Growth | *Reference* |  |  |
| Endpoint category -Mortality | 0.56 | 0.18 | 0.24 – 0.94 |
| Endpoint category -Reproduction | 0.46 | 0.22 | 0.05 – 0.93 |
| Exposure duration | 0.02 | 0.01 | -0.01 – 0.04 |
| Material type × Particle shape –SS Irregular | *Reference* |  |  |
| Material type × Particle shape –MP Irregular | -1.07 | 0.71 | -2.56 – 0.27 |
| Material type × Particle shape –MP Spherical | -0.70 | 0.52 | -1.72 – 0.37 |
| Grain size class | 0.15 | 0.16 | -0.11 – 0.52 |
| **Monotonic effects** |  |  |  |
| Grain size class -1 | 0.39 | 0.24 | 0.02 – 0.83 |
| Grain size class -2 | 0.17 | 0.16 | 0 – 0.61 |
| Grain size class -3 | 0.21 | 0.17 | 0.01 – 0.63 |
| Grain size class -4 | 0.23 | 0.18 | 0.01 – 0.67 |
| **Random effects** |  |  |  |
| Study | 1.44 | 0.34 | 0.85 – 2.14 |
| Study:Species | 0.62 | 0.23 | 0.27 – 1.17 |
| Residual | 0.31 | 0.08 | 0.17 – 0.47 |

Table S2. Description of parameters used for the standardization models where model 1 is the precursory standardization model used as input to the pSSD+ model and model 2 is the stand-alone Bayesian mixed model used to compare the toxicity of microplastics and suspended sediments.

| Model | Response | Expl. variable | Variable type | Meaning | Unit/Levels | Prediction value |
| --- | --- | --- | --- | --- | --- | --- |
| Model 1 | Toxicity value (mm^3^/L) | Duration days | Continuous | Exposure duration | Days | 28 |
|  |  | Material type class x Particle shape | Factor/Interaction | Plastic or mineral material combined with typical particle shape (irregular or spherical). Spherical only for plastic. | Irregular MP or SS or spherical MP | MP- or SS-irregular |
|  |  | Grain size class^1^ | Ordered factor | Size class according to sediment grain classes | Clay, silt, fine sand, sand etc. | Clay |
| Model 2 | eNOEC (mm^3^/L )* | Duration days | Continuous | Exposure duration | Days |  |
|  |  | Particle exposure | Factor | Natural environment for a species and lifestage | Turbid, average or clear |  |
|  |  | Material type class x Particle shape | Factor/Interaction | Plastic or mineral material combined with typical particle shape -irregular or spherical. Spherical only for plastic. | MP or SS -Irregular or Spherical shape |  |
|  |  | Endpoint category | Factor | Endpoint class reflecting a higher order effect. | Mortality, growth, reproduction |  |
|  |  | Species class | Factor | Organism type | e.g. shrimp, cladoceran, bivalve etc. |  |
|  |  | Grain size class^1^ | Ordered factor | Size class according to sediment grain classes | Clay, silt, fine sand, sand etc. |  |
|  |  | Study^2^ | Factor |  |  |  |
|  |  | Study:Species^2^ | Factor/Interaction | Species | e.g. *Daphnia magna*, *Mytilus edulis within a particular study* |  |

Note:

^1^ Grain size class was modeled as a monotonic effect (Bürkner 2017),

^2^ Uncertainty associated with variability between studies and species was accounted for by inclusion of the variables *Study* and *Species* as factors with random intercepts. The estimated NOEC (eNOEC) was log10-transformed to stabilize the model residuals.

*Choice of priors*

We used weakly informative normal priors for all β-parameters in both Bayesian models since the default flat priors have been shown to be unnecessarily conservative (Lemoine 2019). The only exception was for the variable *Grain size class* which was modelled as a monotonic effect and hence had a Dirchlet prior for the simplex monotonic parameter. The Bayesian mixed model which was parameterized with Student-t errors used a Gamma distributed prior for the ν parameter, i.e. the degrees of freedom for the Student-t distribution. The intercept was Student-t distributed and the standard deviations for the random effects followed a Cauchy distribution (Lemoine 2019, McElreath 2020). Details on the prior specifications are provided in tables S3 and S5.

*Table S3. Prior specifications for the precursory pSSD+-standardization model.*

| Prior | Class | Coef |
| --- | --- | --- |
| Normal (0,10) | b | Exposure duration |
| Normal (0,10) | b | Feeding strategy, Filter feeder |
| Normal (0,10) | b | Feeding strategy, Predator |
| Normal (0,10) | b | Material type:Particle shape, MP Irregular |
| Normal (0,10) | b | Material type:Particle shape, MP Spherical |
| Normal (0,10) | b | Grain size class |
| Normal (0,10) | b | Particle shape, Spherical |
| Normal (0,10) | b | Species class, Artemia |
| Normal (0,10) | b | Species class, Bivalve |
| Normal (0,10) | b | Species class, Calanoid copepod |
| Normal (0,10) | b | Species class, Cladoceran |
| Normal (0,10) | b | Species class, Crab |
| Normal (0,10) | b | Species class, Cyclopoid copepod |
| Normal (0,10) | b | Species class, Fairy shrimp |
| Normal (0,10) | b | Species class, Fish |
| Normal (0,10) | b | Species class, Gammarid |
| Normal (0,10) | b | Species class, Insect |
| Normal (0,10) | b | Species class, Krill |
| Normal (0,10) | b | Species class, Mysid |
| Normal (0,10) | b | Species class, Polychaete |
| Normal (0,10) | b | Species class, Rotifer |
| Normal (0,10) | b | Species class, Shrimp |
| Normal (0,10) | b | Species class, Tunicate |
| Normal (0,10) | b | Species class, Urchin |
| student-t (3 ,0 ,2.5) | Intercept |  |
| student-t (3 ,0 ,2.5) | sigma |  |
| dirichlet(1) | simo | Grain size class |

Table S4. Uncertainty factors used to convert dose descriptors into NOECs. Modified from Wigger et al. (2020).

| **Type of dose descriptor** | **UF_dose_ modal value** | **Reference** |
| --- | --- | --- |
| LC_25-50,_ EC_25-50,_ IC_25-50_ | 10 | Kim *et al.* 2011, Schwab *et al.* 2011 |
| LC_20,_ EC_20,_ IC_20_ | 2 | Azimonti et al. 2015, Beasley et al. 2015 |
| LOEC | 2 | ECHA 2008 |
| LC_10,_ EC_10,_ IC_10_ | 1 | Azimonti et al. 2015, Beasley et al. 2015 |
| HONEC | 1 | Equal to NOEC |

Note: HONEC = highest observed no-effect concentration, LOEC = lowest observed effect concentration, NOEC = no observed effect concentration, IC_x_ = Inhibitory concentration, EC_x_ = effect concentration, LC_x_ = lethal concentration, UF_dose_ = uncertainty factor for a dose descriptor.

Table S5. Prior specifications for the Bayesian mixed model

| Prior | Class | Coef |
| --- | --- | --- |
| Normal (0,10) | b | Exposure duration |
| Normal (0,10) | b | Endpoint category, Mortality |
| Normal (0,10) | b | Endpoint category, Reproduction |
| Normal (0,10) | b | Feeding strategy, Filter feeder |
| Novrmal (0,10) | b | Feeding strategy, Predator |
| Normal (0,10) | b | Material type:Particle shape, MP Irregular |
| Normal (0,10) | b | Material type:Particle shape, MP Spherical |
| Normal (0,10) | b | Grain size class |
| Normal (0,10) | b | Particle exposure, Low |
| Normal (0,10) | b | Particle exposure, Medium |
| Normal (0,10) | b | Particle shape, Spherical |
| student-t (3, 0, 2.5) | Intercept |  |
| gamma (2, 0.1) | nu |  |
| cauchy (0, 1) | sd |  |
| cauchy (0, 1) | sd | Study |
| cauchy (0, 1) | sd | Intercept, Study |
| cauchy (0, 1) | sd | Study:Species |
| cauchy (0, 1) | sd | Intercept, Study:Species |
| cauchy (0, 1) | sigma |  |
| dirichlet(1) | simo | Grain size class |

Table S6. Descriptive statistics for variables in the selected data that are relevant to exposure conditions. Grain size classes are coded 1-5 (clay, silt, very fine sand, fine sand, medium sand).

| Particle type | Variable | n | mean | sd | median | min | max | range | skew | kurtosis | se |
| --- | --- | --- | --- | --- | --- | --- | --- | --- | --- | --- | --- |
| Suspended sediments | Exposure duration (days) | 77 | 10.87 | 12.78 | 8.00 | 0.04 | 71.00 | 70.96 | 2.94 | 10.43 | 1.46 |
|  | Particle density (g/cm^3^) | 77 | 2.57 | 0.14 | 2.60 | 2.00 | 2.65 | 0.65 | -3.41 | 10.6 | 0.02 |
|  | Grain size class (1-5) | 77 | 1.16 | 0.37 | 1.00 | 1.00 | 2.00 | 1.00 | 1.86 | 1.48 | 0.04 |
|  | Highest test conc. (mm^3^/L) | 77 | 6.67 × 10^3^ | 14.73 x 10^4^ | 308.08 | 1.34 | 4.50 × 10^4^ | 4.50 × 10^5^ | 2.02 | 2.2 | 1.68 × 10^3^ |
|  | Lowest test conc. (mm^3^/L) | 62 | 648.99 | 4.88 × 10^3^ | 20.19 | 4.46 × 10^-3^ | 3.85 × 10^5^ | 3.85 × 10^5^ | 7.5 | 55.09 | 619.91 |
| Microplastic | Exposure duration (days) | 123 | 13.16 | 17.15 | 4.00 | 0.13 | 60.00 | 59.87 | 1.74 | 2.16 | 1.55 |
|  | Particle density (g/cm^3^) | 123 | 1.04 | 0.13 | 1.05 | 0.92 | 1.40 | 0.48 | 1.43 | 0.94 | 0.01 |
|  | Grain size class (1-5) | 123 | 2.53 | 1.20 | 2.00 | 1.00 | 5.00 | 4.00 | 0.37 | -0.94 | 0.11 |
|  | Highest test conc. (mm^3^/L) | 123 | 103.51 | 242.94 | 11.04 | 6.50 × 10^-4^ | 1.54 × 10^3^ | 1.54 × 10^3^ | 3.75 | 14.33 | 21.9 |
|  | Lowest test conc. (mm^3^/L) | 123 | 35.93 | 51.20 | 0.20 | 2.38 × 10^-4^ | 176.99 | 176.99 | 0.89 | -0.88 | 4.62 |

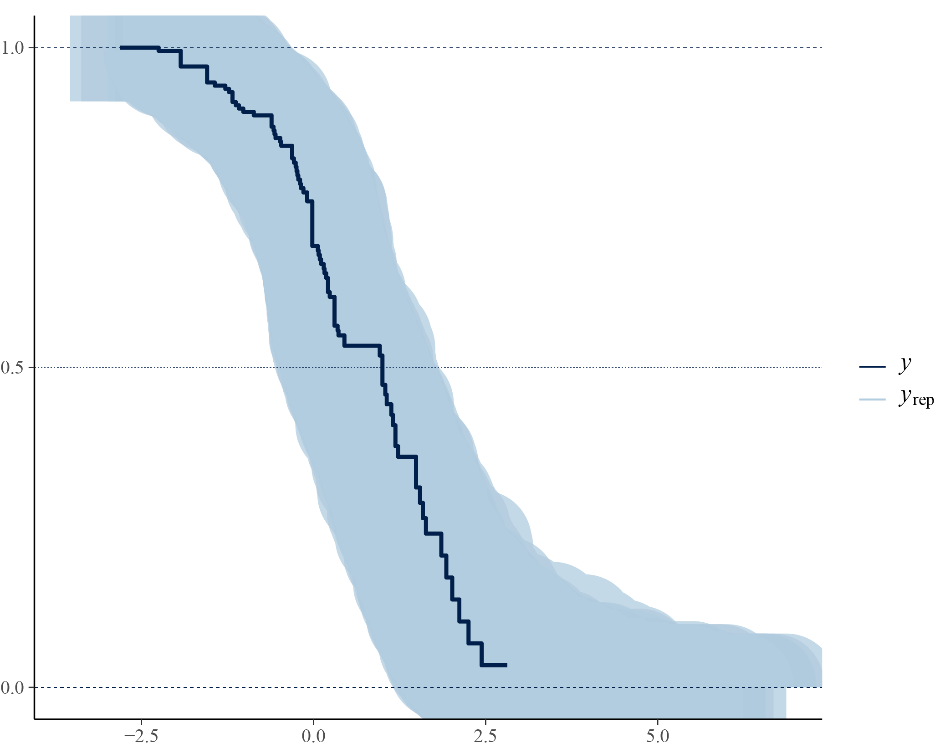

Figure S1. Simulated residual plot for the Bayesian mixed model (Table S2, model 2). Dark line (y) shows empirical data and light blue, shaded area (y_rep_) the simulated residuals.

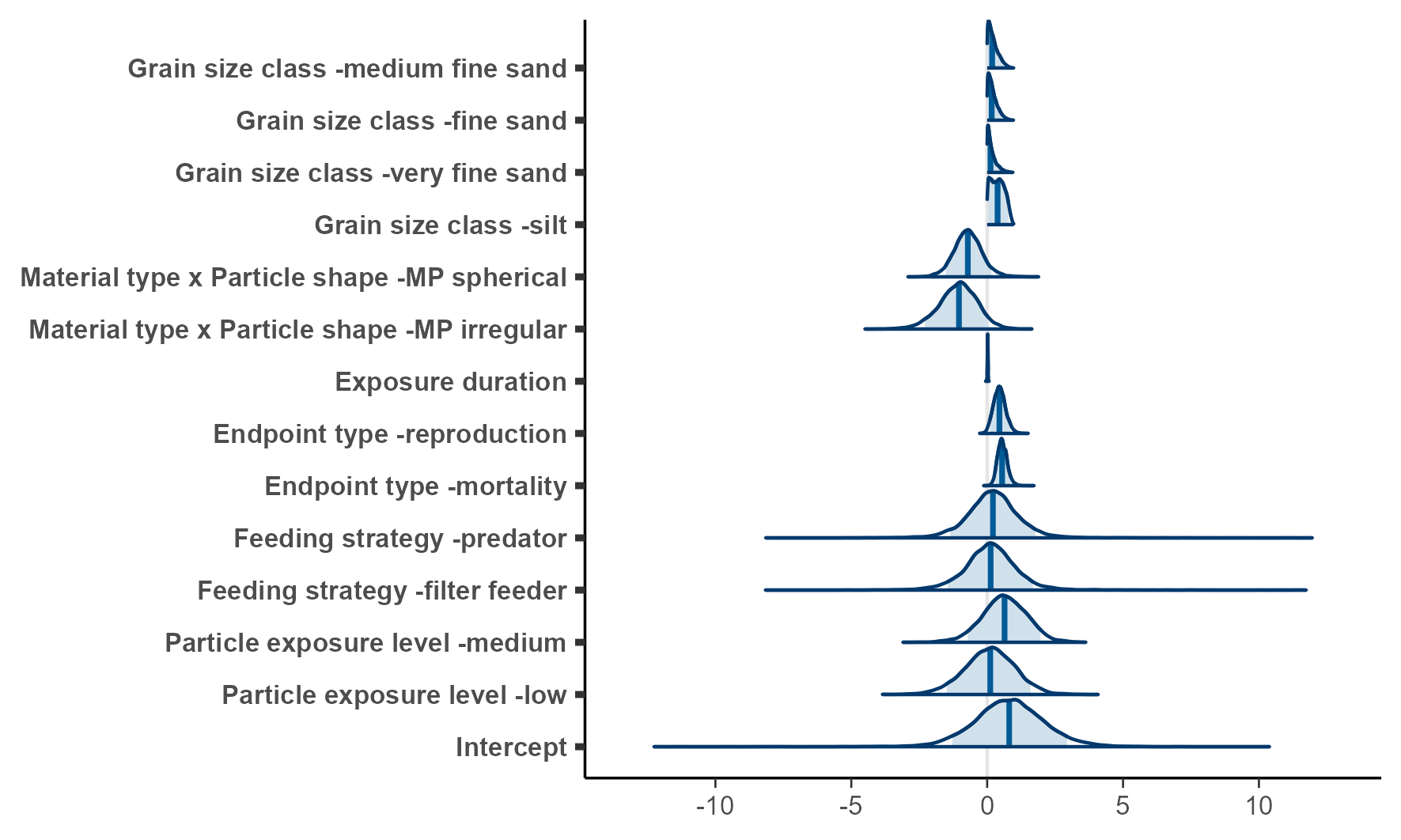

Figure S2. Posterior distribution of regression coefficients from the Bayesian mixed model. The density plots are scaled to equal height for visibility.

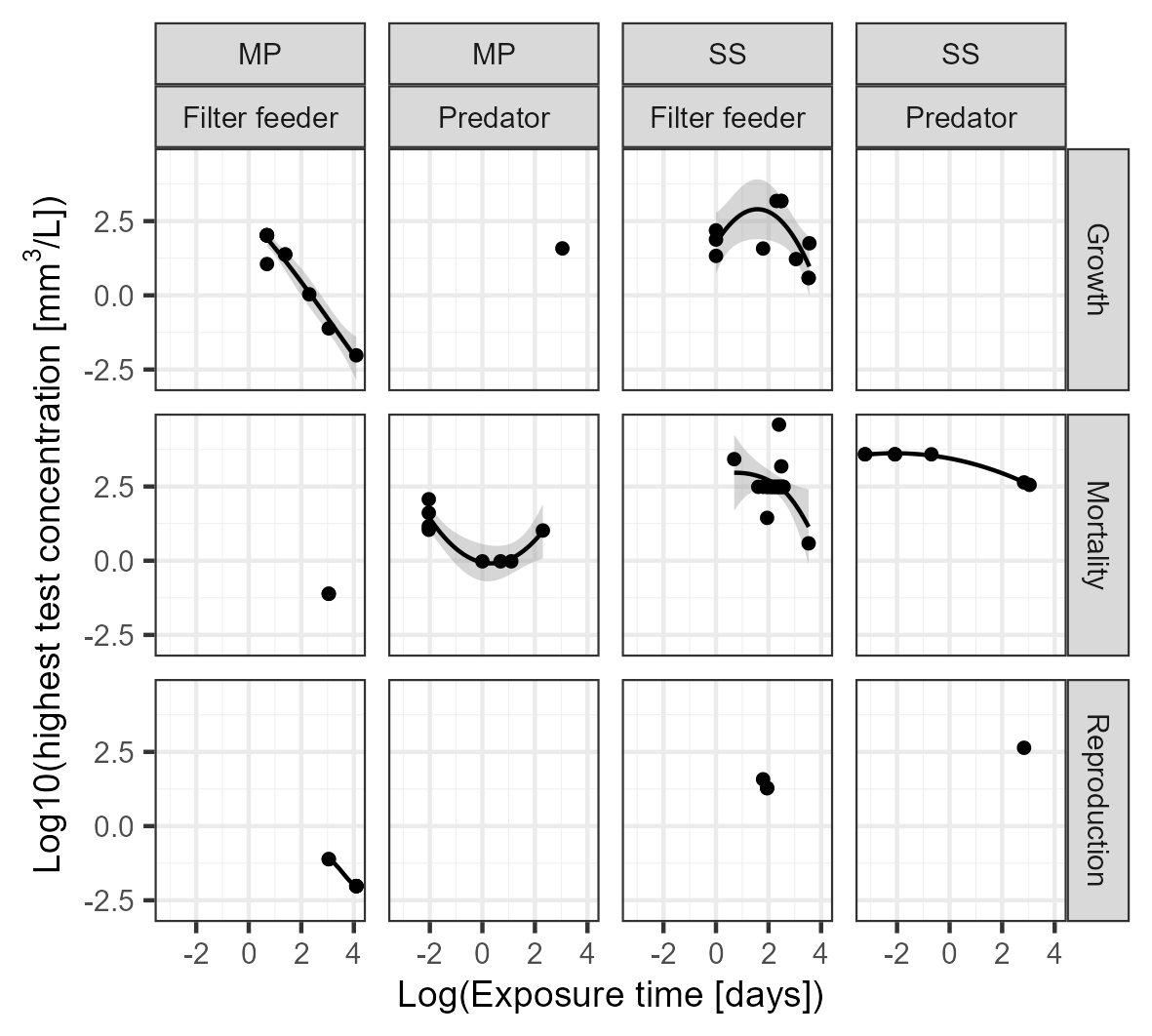

Figure S3. Log10-transformed highest used test concentration as a function of Log-transformed exposure time divided per particle type (microplastic, MP vs suspended sediments, SS), feeding type and endpoint.

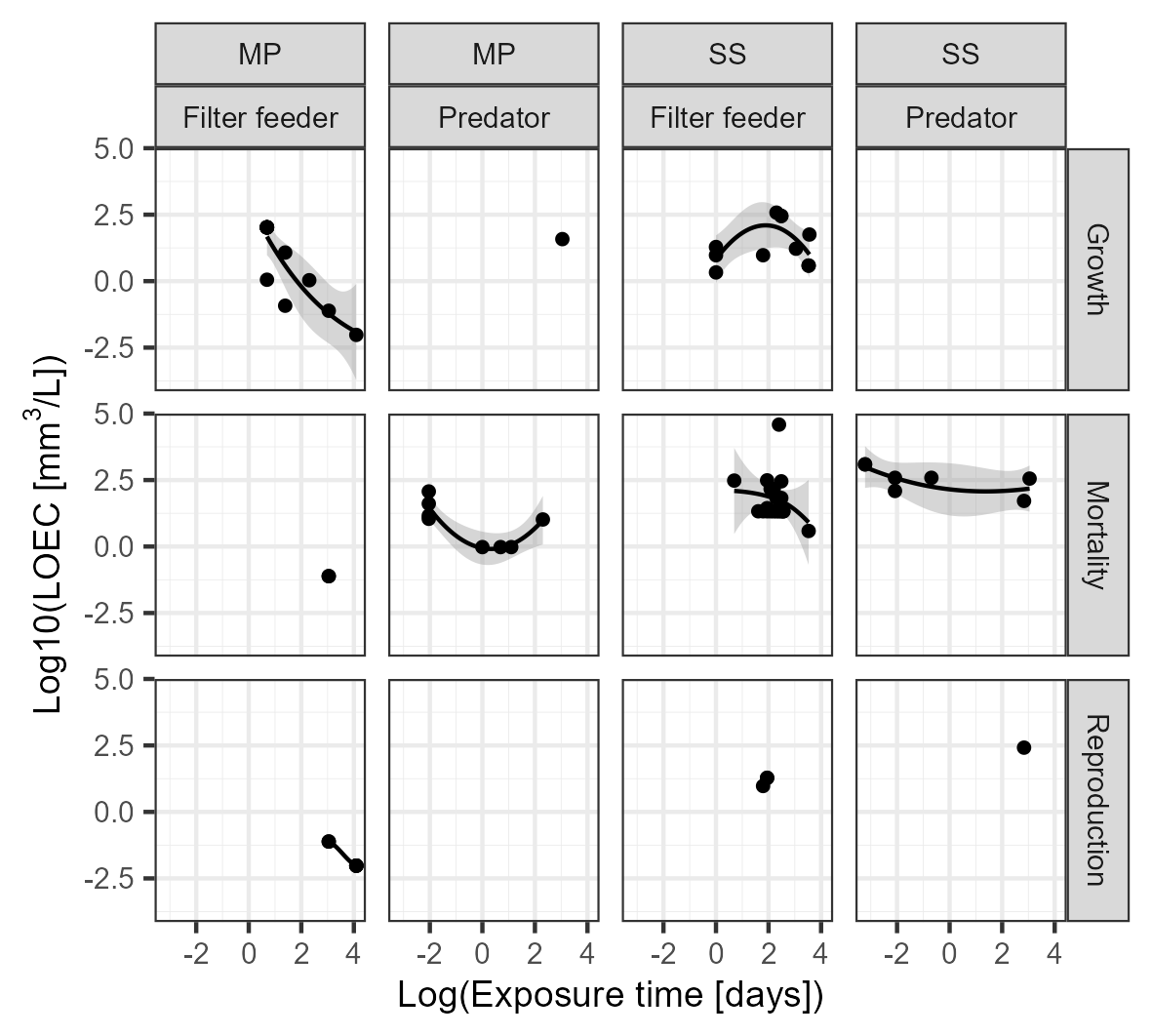

Figure S4. Log10-transformed Lowest Effect Concentration (LOEC) as a function of Log-transformed exposure time divided per particle type (microplastic, MP vs suspended sediments, SS), feeding type and endpoint.

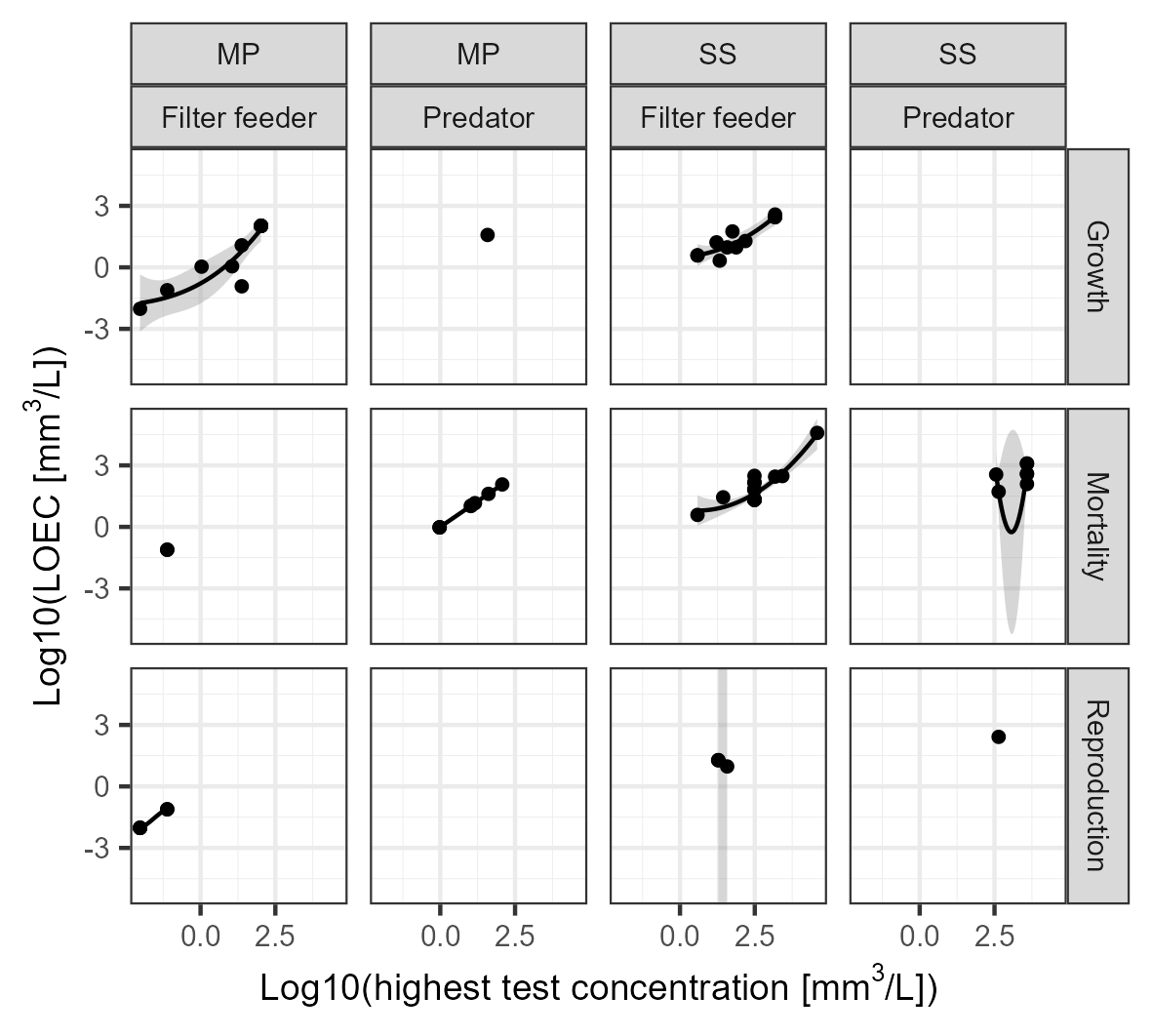

Figure S5. Log10-transformed Lowest Effect Concentration (LOEC) as a function of Log10-transformed highest used test concentration divided per particle type (microplastic, MP vs suspended sediments, SS), feeding type and endpoint.

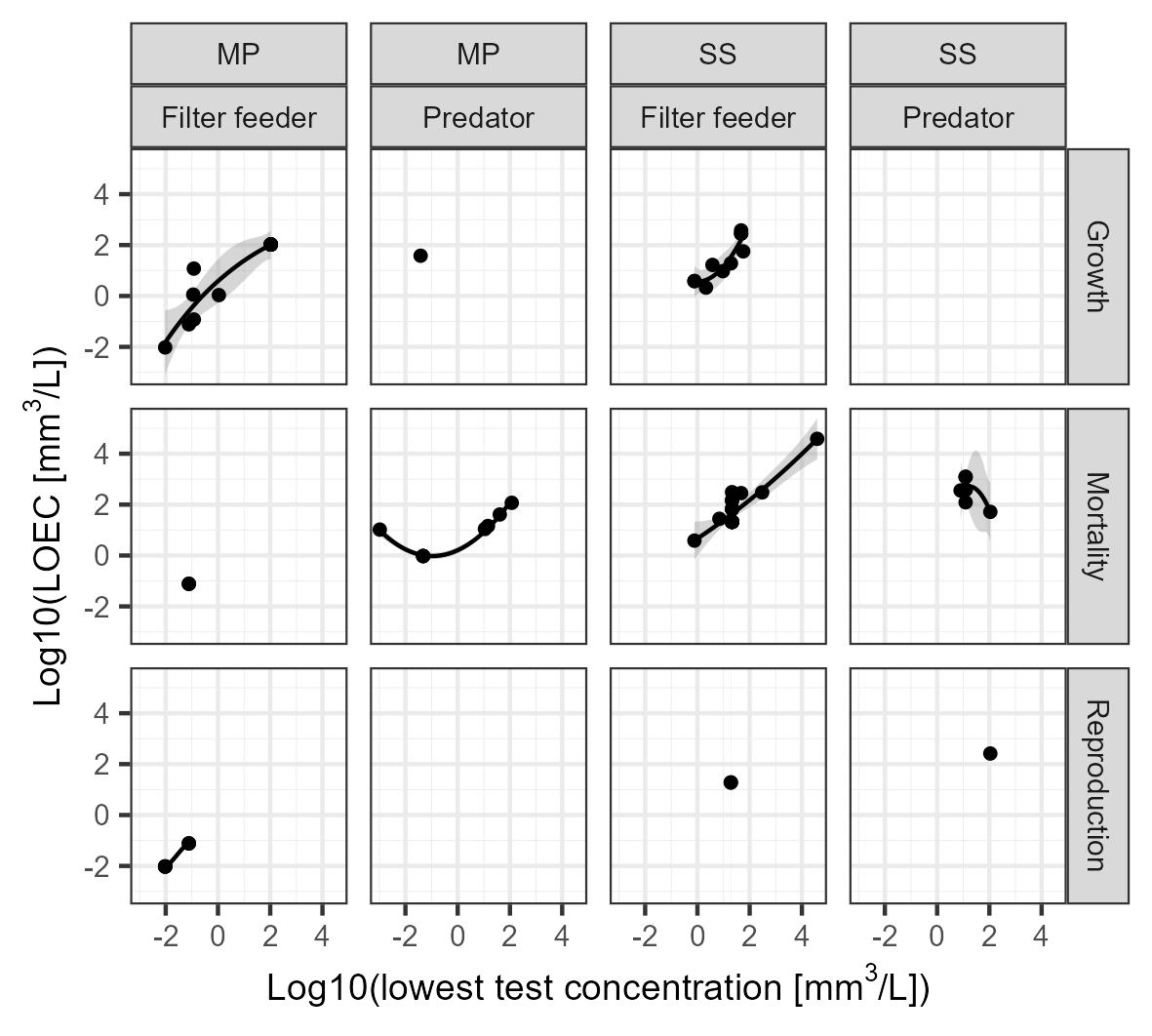

Figure S6. Log10-transformed Lowest Effect Concentration (LOEC) as a function of Log10-transformed lowest used test concentration (zeros excluded) divided per particle type (microplastic, MP vs suspended sediments, SS), feeding type and endpoint.

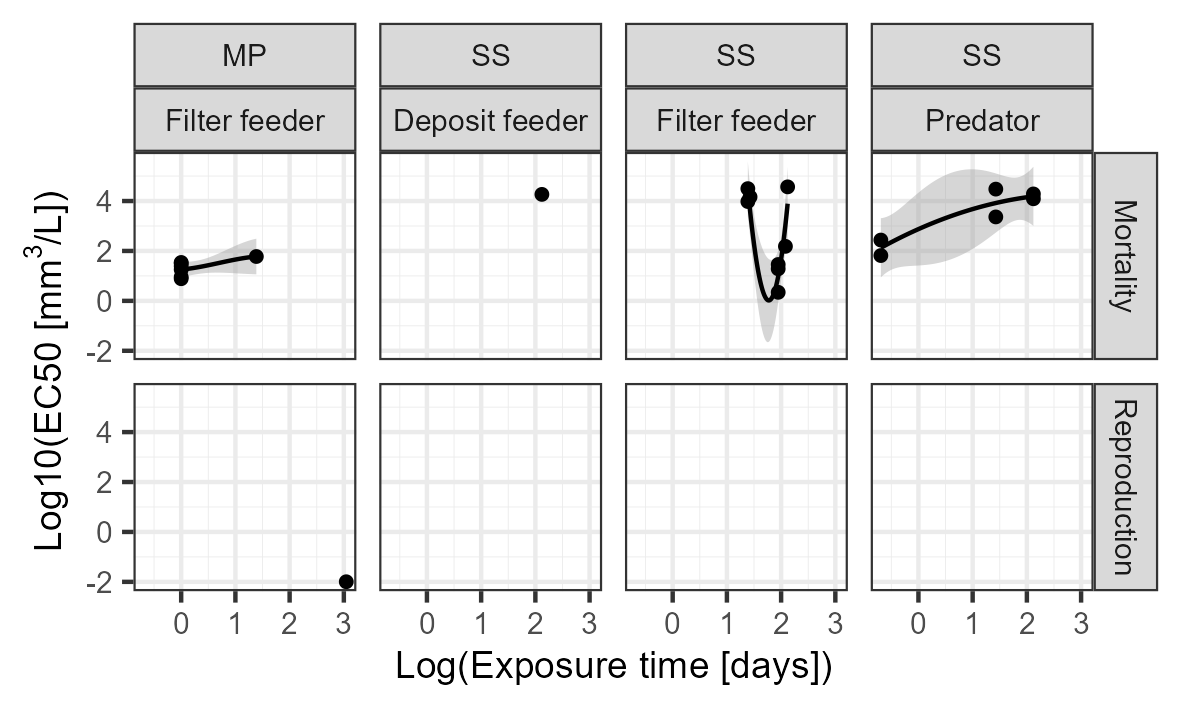

Figure S7. Log10-transformed EC_50_-values as a function of Log-transformed exposure time divided per particle type (microplastic, MP vs suspended sediments, SS), feeding type and endpoint.
